# T-cells specific for KSHV and HIV migrate to Kaposi sarcoma tumors and persist over time

**DOI:** 10.1101/2024.02.06.579223

**Authors:** Shashidhar Ravishankar, Andrea M.H. Towlerton, Iyabode L. Tiamiyu, Peter Mooka, Janet Nankoma, James Kafeero, Dennis Mubiru, Semei Sekitene, Lauri D. Aicher, Chris P. Miller, David G. Coffey, Lazarus Okoche, Prisca Atwinirembabazi, Joseph Okonye, Jessica White, David M. Koelle, Lichen Jing, Warren T. Phipps, Edus H. Warren

## Abstract

Kaposi sarcoma-associated herpesvirus (KSHV) is the etiologic agent of Kaposi sarcoma (KS), which causes significant morbidity and mortality worldwide, particularly in people living with HIV (PLWH) and in sub-Saharan Africa where KSHV seroprevalence is high. Postulating that T-cells specific for KSHV and HIV would be attracted to KS tumors, we performed transcriptional profiling and T-cell receptor (TCR) repertoire analysis of tumor biopsies from 144 Ugandan adults with KS, 106 of whom were also living with HIV. We show that CD8**^+^** T-cells and M2-polarized macrophages are the most common immune cells in KS tumors. The TCR repertoire of T-cells associated with KS tumors is shared across spatially and temporally distinct tumors from the same individual. Clusters of T-cells with predicted shared specificity for uncharacterized antigens, potentially encoded by KSHV or HIV, comprise ∼25% of the T-cells in KS tumors. Single-cell RNA-sequencing of blood from a subset of 9 adults captured 4,283 unique αβ TCRs carried in 14,698 putative KSHV- or HIV-specific T-cells, which carried an antigen-experienced effector phenotype. T-cells engineered to express a representative sample of these TCRs showed high-avidity recognition of KSHV- or HIV-encoded antigens. These results suggest that a polyspecific, high-avidity KSHV- and HIV-specific T-cell response, potentially inhibited by M2 macrophages, migrates to and localizes with KS tumors. Further analysis of KSHV- and HIV-specific T-cells in KS tumors will provide insight into the pathogenesis of KS and could guide the development of specific immune therapy based on adoptive transfer or vaccination.

**Author Summary:** In this work, we set out to examine Kaposi Sarcoma (KS) tumor tissue, as well as peripheral blood cells from individuals with KS, both living with and without HIV. Our goal was to identify T-cells that specifically recognize antigens encoded by Kaposi sarcoma-associated herpesvirus (KSHV) or HIV. By analyzing the T-cell repertoire in KS tumor biopsies from people in Uganda with different types of KS, we uncovered clusters of T-cells with previously unknown ability to recognize these viruses. Through single-cell sequencing of peripheral blood cells, we also observed that many of these T-cells had cell-killing properties. Notably, they often coexisted with a subset of macrophages with immunosuppressive properties, which we suspect may be suppressing the function of virus-targeting T-cells. Our findings suggest that additional studies of these virus-targeting T-cells and their interaction with immunosuppressive macrophages could significantly advance the development of effective therapeutics against KS.

## Introduction

Kaposi sarcoma-associated herpesvirus (KSHV) is the etiologic agent of Kaposi sarcoma (KS), primary effusion lymphoma, and multicentric Castleman’s disease. KS causes significant morbidity and mortality worldwide, particularly in people living with HIV (PLWH) and in sub-Saharan Africa (SSA) where KSHV seroprevalence is high. It is estimated that 80% of the KS burden in SSA, where the impact of KS is heaviest, is attributable to HIV infection (1). KS most often develops in the setting of T-cell deficiency or dysfunction, such as in KSHV-seropositive individuals with HIV infection or KSHV-seropositive recipients of solid organ or allogeneic hematopoietic cell transplants. In these settings KS can remit following initiation of antiretroviral therapy (ART) or withdrawal of immune suppression. In SSA, primary infection with KSHV is thought to occur in childhood, but most cases of KS and other KSHV-associated disease in both PLWH and people without HIV infection develop many years, often several decades, later. These observations suggest that loss or impairment of a T-cell component of pre-existing KSHV-specific immunity underlie the development of these diseases. Strategies that preserve or restore the T-cell component of KSHV-specific immunity in PLWH and others at risk should, therefore, have potential for the prevention or treatment of KSHV-associated disease.

Transcriptional profiling of KS tumors by our group (2) and others (3, 4) has demonstrated that they contain high levels of KSHV transcripts. We therefore postulated that KS tumors should attract significant numbers of T-cells that are specific for KSHV-encoded antigens. Moreover, we speculated that KS tumors in PLWH might also contain HIV transcripts and attract HIV-specific T-cells. To identify T-cells specific for KSHV or HIV, we used adaptive immune receptor repertoire sequencing (AIRR-seq) to define and enumerate the T-cell receptors (TCRs) expressed in 299 KS tumor biopsies and 64 samples of uninvolved axillary skin from 106 PLWH with KS (epidemic KS) and 38 HIV-seronegative adults with KS (endemic KS). For many individuals, it was possible to analyze multiple spatially distinct tumors obtained at multiple time points over one year of observation. To further characterize the KS TME, we profiled the transcriptome of 51 KS tumors (39 from PLWH (2) and 12 from HIV-seronegative adults) using bulk RNA sequencing (RNA-seq) and performed targeted gene expression analysis on 27 additional tumors and 5 samples of normal uninvolved skin. Finally, we performed single-cell RNA-seq of 20 PBMC samples from nine of the 144 individuals, 5 of whom were living with HIV, to capture TCR αβ pairs carried in putative KSHV- or HIV-specific T-cells, and we investigated the KSHV- or HIV-specificity of a sample of these TCRs.

Our results demonstrate that CD8**^+^** T-cells and macrophages with M2 polarization are the most common immune cell types in KS tumors. Analysis of the TCR repertoire in KS tumors reveals a polyclonal idiosyncratic T-cell response that is largely private to each individual but shared across spatially and temporally distinct tumors from the same individual. Clustering TCR β chain CDR3 amino acid sequences by sequence similarity, we find that ∼25% of the TCRs expressed by T-cells in KS tumor biopsies fall into antigenic specificity groups with strong MHC association that are predicted to recognize similar or identical peptide/MHC complexes. We identified 14,698 T-cells bearing 4,283 unique αβ TCRs that belonged to antigenic specificity groups with predicted specificity for KSHV or HIV and characterized the phenotype of these T-cells. Finally, we confirmed the specificity for KSHV- or HIV-encoded peptides of a representative subset of these TCRs.

## Results

### KSHV and HIV gene expression in KS tumors

RNA-seq of 39 epidemic (HIV-associated) and 12 endemic (HIV-negative) KS tumors revealed consistent expression of both latent and lytic cycle KSHV genes within the TME. Kaposin (*K12*) showed the highest expression of all KSHV genes in all but two tumors (Figure 1A) and was also the most strongly expressed gene in the 22 tumors evaluated with the targeted expression assay (Supplementary Figure 1).

**Figure 1:**
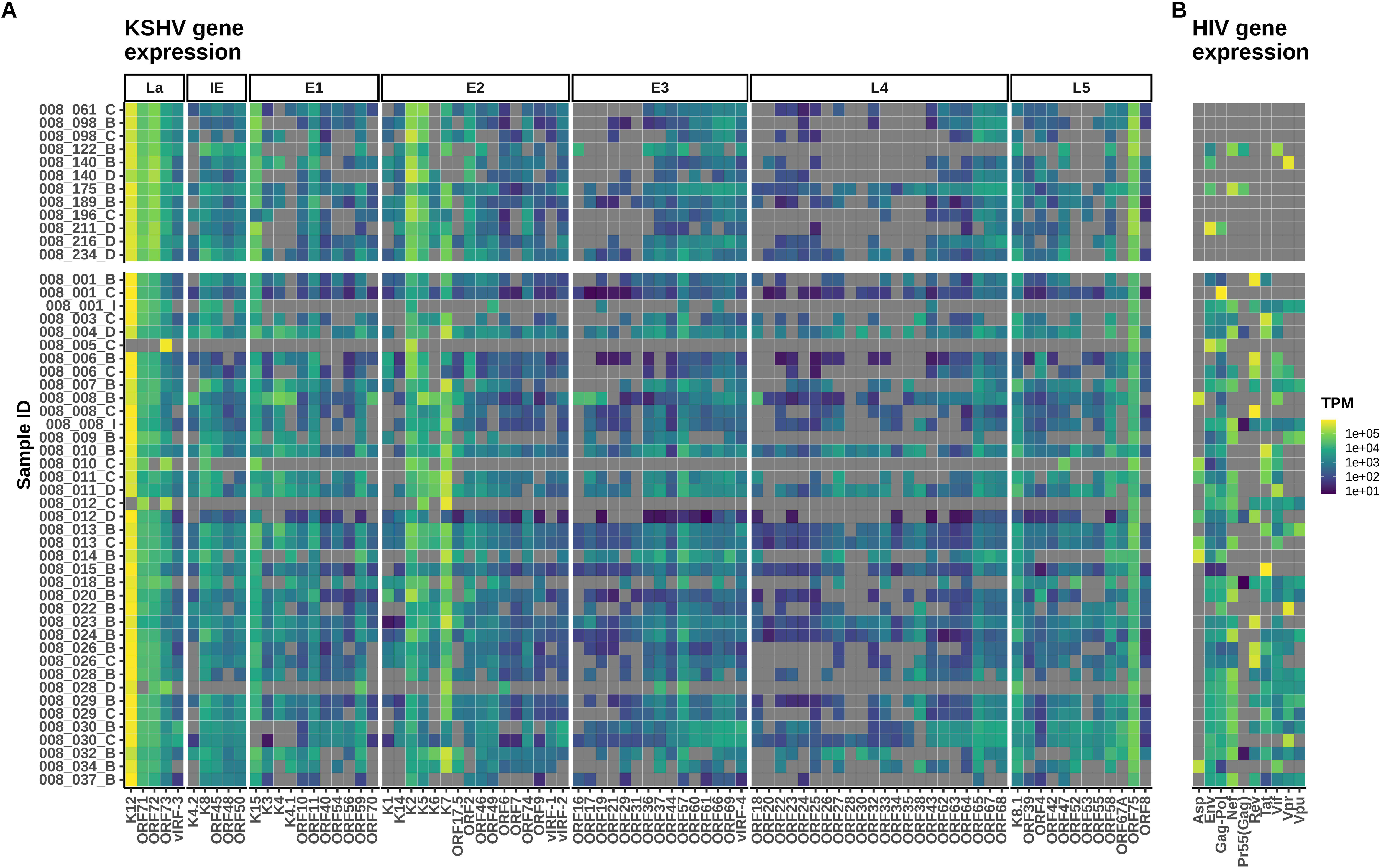
RNA-seq confirms expression of KSHV and HIV genes in KS tumors. (A) KSHV gene expression in 12 endemic (top) and 39 epidemic (bottom) KS tumors. Latent (La), Immediate early (IE), Early lytic stage 1 (E1), Early lytic stage 2 (E2), Early lytic stage 3 (E3), Late lytic stage 4 (L4), and Late lytic stage 5 (L5). (B) Expression of HIV genes across all 51 KS tumors.

The key latency gene *ORF73* (LANA-1) and the key lytic gene *ORF75* were both highly expressed across all tumor samples, reflecting the dynamic state of KSHV within the KS TME. Consistent expression of other latency genes (*e.g.*, *ORF71, ORF72*) as well as early lytic genes (*e.g.*, *K15*, *K2*, *K7*) was also observed. HIV gene expression was detected in all 39 tumors obtained from individuals with epidemic KS and 4 of the 12 tumors obtained from individuals with endemic KS, all of whom were ART-naive at study entry (Figure 1B). HIV Gag/Pol was expressed in all 39 samples with epidemic KS, suggesting active replication of HIV within the TME.

### M2 macrophages and CD8^+^ T-cells are the dominant cell types in KS tumors

Deconvolution of the RNA-seq data from KS tumors using CIBERSORTx (5) was performed to estimate the relative size of immune cell subsets in the KS TME. Macrophages with M2 polarization were predicted to be the most common immune cell type, accounting for a mean of 38% of the immune cells in endemic KS tumors and 22% of the immune cells in epidemic KS tumors (Figure 2A, 2B).

**Figure 2:**
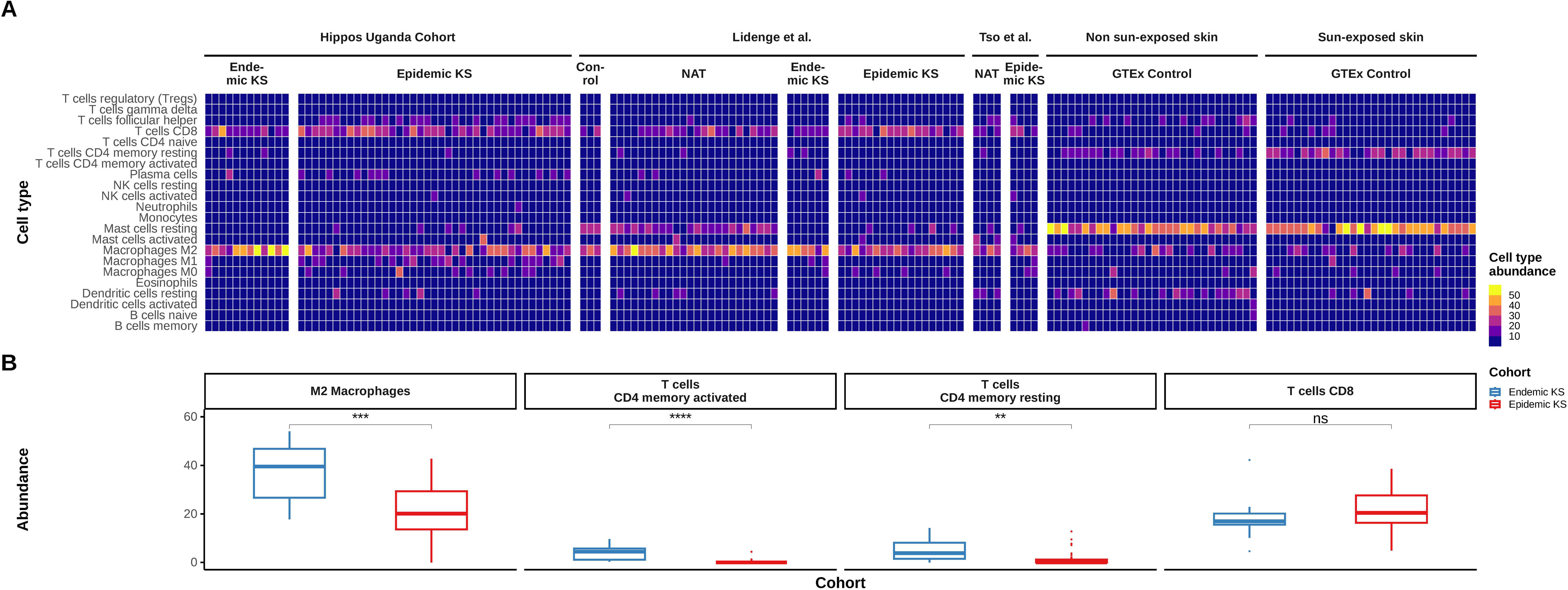
KS tumors show an enrichment of M2 macrophages and CD8^+^ T-cells in comparison to control skin samples. (A) Heatmap of relative abundance of the indicated immune cell types found in KS tumors and skin samples from individuals with KS as well as control skin samples from the GTEx consortium, determined via deconvolution of RNA-seq data using CIBERSORTx. The RNA-seq datasets analyzed were derived from (left to right) 12 endemic and 39 epidemic KS tumors from the HIPPOS cohort from Uganda, 3 control skin samples, 24 paired NAT and KS tumor samples from 6 individuals with endemic KS and 18 individuals with epidemic KS from (4); 4 paired NAT and tumor samples from individuals with epidemic KS from (3); and 30 non-sun-exposed control skin samples and 30 sun-exposed control skin samples from the GTEx consortium (6). (B) Box and whisker plots showing the mean and interquartile range (IQR) of the percentages of M2 macrophages, activated CD4**^+^** memory T-cells, resting CD4**^+^** memory T-cells, and CD8**^+^**T-cells, as estimated using CIBERSORTx, in the endemic (blue) and epidemic (red) KS tumors in (A). The upper/lower whiskers extend from the hinge to the largest/smallest value, respectively, no further than 1.5 * IQR from the hinge. Significance between comparisons is indicated by ns (p > 0.05), ** (p<= 0.01), *** (p<= 0.001), **** (p <= 0.0001).

The higher proportion of M2 macrophages in endemic compared with epidemic KS tumors was statistically significant. CD8**^+^** T-cells were predicted to be the second largest group of immune cells within the TME, accounting for a mean of 18% and 21% of the cells in the endemic and epidemic tumors, respectively. CD4**^+^** resting memory T-cells and CD4**^+^** activated memory T-cells were predicted to make up 5% and 4%, respectively, of the immune cells in endemic KS tumors but only 1.3% and 0.2% of the immune cells in epidemic KS tumors (Figure 2A, 2B). Cell type deconvolution of previously published RNA-seq datasets (3, 4) from 28 KS tumor – NAT tissue pairs and skin from 3 non-KS control subjects similarly revealed that M2-polarized macrophages and CD8**^+^** T-cells were the most common immune cells in the KS tumors (Figure 2A). In contrast, deconvolution analysis of RNA-seq data from normal sun-exposed and non-sun-exposed skin from the GTEx consortium (6) showed a predominance of resting mast cells but low levels of M2 macrophages and few, if any, CD8**^+^** T-cells (Figure 2A).

We observed a negative correlation between the relative abundance of M2 macrophages and CD8**^+^**T-cells across all 51 KS tumors from this study as well as the 28 previously published KS tumor datasets (Supplementary Figure 2).

We observed a stronger negative correlation in endemic KS tumors (this study: R = -0.61, p = 0.035; Lidenge *et al.* (4),: R= -0.88, p = 0.019) compared to epidemic KS tumors (this study: R = -0.23, p = 0.162; Tso *et al.*: R = -0.82, p = 0.18; Lidenge *et al.*: R = -0.59; p = 0.01), with the exception of the 4 KS tumors from Tso *et al.* (3) (Supplementary Figure 2). A similar negative correlation was observed across NAT samples from individuals with epidemic KS (Lidenge *et al.:* R = -0.19; p = 0.44; Tso *et al.*: R = -0.96, p = 0.04). However, in the limited number of NAT samples from individuals with endemic KS, there was a weak positive correlation between abundance of M2 macrophages and CD8**^+^** T-cells (Lidenge *et al.:* R = 0.17, p = 0.74).

### TCR repertoire diversity is higher in endemic than in epidemic KS tumors

We performed adaptive immune repertoire sequencing (AIRR-seq) of the TCR β chain repertoire in KS tumor biopsies obtained at the time of study enrollment from all 144 individuals in our Uganda KS cohort. This effort generated 2.85 million TCR β chain sequences. AIRR-seq was also performed on NAT samples from uninvolved axillary skin from 65 of the 144 individuals. For comparison we analyzed the TCR β chain repertoire in tumor biopsies from 49 children and adolescents with Epstein-Barr virus-associated Burkitt lymphoma (BL) from Uganda and Ghana (7). We used the generalized entropy measure, Rényi entropy, calculated at different values of the parameter α (alpha), to assess the diversity of the TCR repertoires at the time of study enrollment across the different groups. T-cell repertoires in endemic KS tumors showed significantly higher species richness (α = 0), Shannon entropy (α = 1), and Simpson’s diversity (α = 2) when compared to epidemic KS tumors and BL tumors from Ghana, but not BL tumors from Uganda (Figure 3A-C).

**Figure 3:**
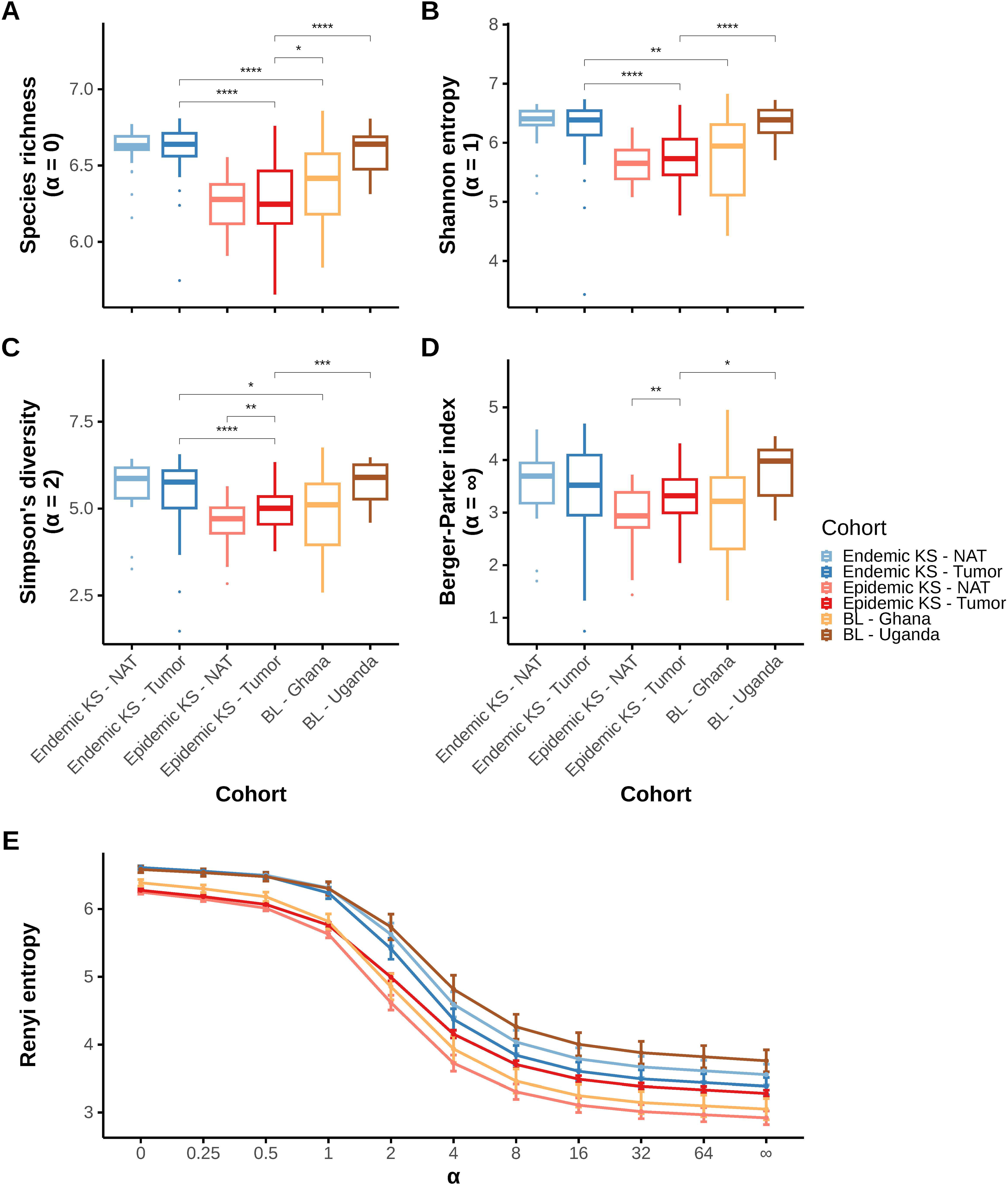
Diversity of infiltrating T-cells is higher in endemic than in epidemic KS tumors. (A, B, C, D, E) Rényi entropy (RE) comparisons, at different values of α, across tissue samples: epidemic KS – NAT (Pink), epidemic KS – tumor (Red), endemic KS – NAT (Light blue), endemic KS – tumor (Blue), Burkitt lymphoma (BL) tumors from Ghana (Yellow), and BL tumors from Uganda (Brown). (A) Species richness (RE α = 0) and (B) Shannon entropy (RE α = 1) are metrics influenced by the number of unique T-cells found in the repertoire. (C) Simpson’s diversity (RE α = 2) and (D) Berger-Parker index (α = ∞) are metrics that are influenced by expanded T-cell clones in the repertoire. (E) Rényi diversity profiles for all six cohorts show the variation in repertoire diversity with change in α. Significance between comparisons is indicated by ns (p > 0.05), * (p <= 0.05), ** (p<= 0.01), *** (p<= 0.001), **** (p <= 0.0001).

The differences in TCR repertoire diversity between the groups for low values of α were not as prominent in the inverse Berger-Parker index (α = ∞), which weighs the more abundant T-cell clones in the repertoire to a greater extent (Figure 3D). The Rényi curves for the TCR repertoires in endemic KS are consistently higher than those for the repertoires from epidemic KS at all values of the Rényi parameter α, indicating greater overall diversity of the TCR repertoires of T-cells infiltrating endemic KS tumors compared with epidemic tumors (Figure 3E).

### KS TIL carry the signature of a polyclonal response to unknown antigens

To determine whether the 2.85 million TCR β chain sequences we observed in KS tumors have previously been associated with a T-cell response to a specific pathogen or tissue antigen or a specific pathological condition, we systematically interrogated two publicly available databases, VDJdb (8–10) and the McPAS-TCR (11) database, for annotated TCR sequences with high confidence of antigenic specificity (Figure 4A).

**Figure 4:**
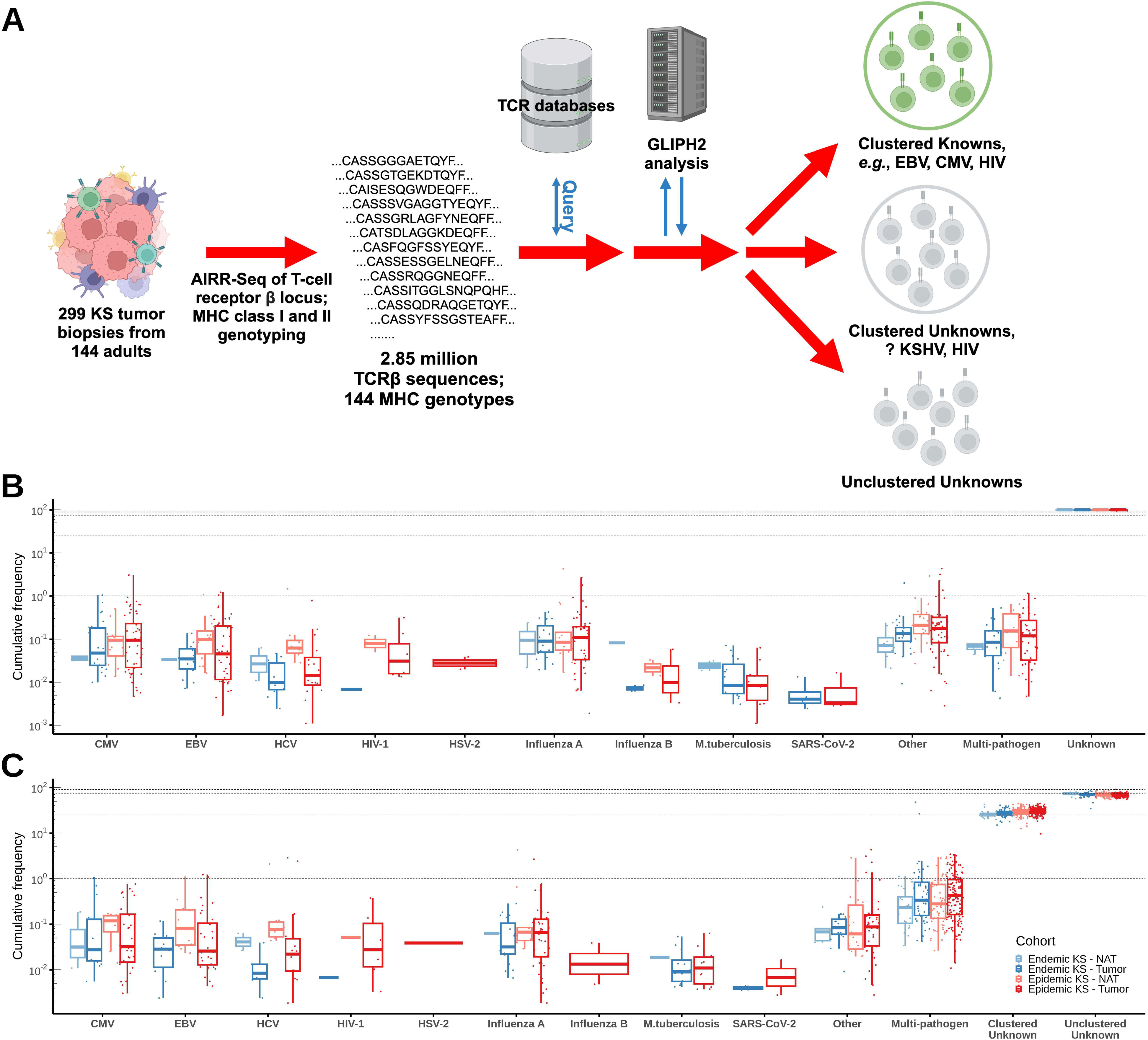
Clusters of T-cells with predicted similar or identical specificity for unknown antigens comprise 25% of the T-cell repertoire in KS tumors. (A) Overview of analytic pipeline of TCR β chain sequences generated from AIRR-seq of 299 KS tumors and from 144 individuals, showing the assignment after GLIPH2 analysis to Clustered Known, Clustered Unknown, or Unclustered Unknown subsets. (B) Mean and interquartile range of the frequency of TCR β chain sequences in the TCR repertoires of individual KS tumors that are listed in VDJdb or the McPAS-TCR database as being associated with a T-cell response to a specific pathogen or tissue antigen, classified according to pathogen and endemic (blue) or epidemic (red) KS. Sequences listed in these databases as being associated with a T-cell response to two or more pathogens are listed as “Multi-pathogen.” (C) Mean and interquartile range of the frequency of TCR β chain sequences assigned by GLIPH2 to antigenic specificity groups associated with T-cell responses to the pathogens in B. The frequency of sequences assigned to Clustered Unknown or Unclustered Unknown antigenic specificity groups is indicated at the far right.

We found that TCR sequences described in VDJdb and/or the McPAS-TCR database and associated with a T-cell response to a specific pathogen or tissue antigen appeared at very low levels (< 1% cumulative frequency) in tumors and NAT samples from individuals in our Uganda cohort with endemic or epidemic KS (Figure 4B). The most frequently associated pathogen/tissue specificities were CMV and Influenza A. The vast majority (99%) of TCR sequences observed in the KS TIL repertoires were not listed in VDJdb or the McPAS-TCR database and thus could not be readily associated with previously defined T-cell responses to specific pathogen or tissue antigens (Figure 4B, far right).

Given the evidence for KSHV and HIV transcripts in KS tumors (Figure 1 and Supplementary Figure 1), we hypothesized that T-cells specific for antigens encoded by KSHV or HIV would be attracted to KS tumors, and that KSHV- and, in HIV**^+^** individuals, HIV-specific TCRs would account for a significant fraction of the TCR repertoire of KS TIL. We used the GLIPH2 algorithm (12) to identify clusters of related TCR β chain sequences in the KS TIL TCR repertoire that were components of TCRs with predicted similar or identical antigenic specificity and MHC restriction (Figure 4A). A small fraction, <1% of the total, of the 45,545 antigenic specificity clusters identified by this analysis contained one or more TCR β chain sequences that are described in VDJdb and/or the McPAS-TCR database as being associated with a T-cell response to a specific pathogen or tissue antigen, and we used the term “Clustered Knowns” to describe these clusters (Figure 4C). However, 24% of the KS TIL T-cell repertoire was encompassed within antigenic specificity groups with as yet unknown specificity, termed “Clustered Unknowns” because they do not contain any sequences listed in public databases as associated with a T-cell response to a defined pathogen or tissue antigen. In total 151,749 unique TCR β chain amino acid sequences were classified as “Clustered Unknowns”. We hypothesize that a significant fraction of the antigenic specificity groups classified as “Clustered Unknowns” are associated with T-cell responses to KSHV, or alternatively to HIV in HIV-seropositive individuals. The remaining 74% of TCR β chain amino acid sequences in tumor and NAT samples from individuals with epidemic and endemic KS (far right in Figure 4C) were not grouped into any specificity groups defined by GLIPH2 and are termed “Unclustered Unknowns.”

### T-cells from “Clustered Unknown” specificity groups migrate to non-contiguous KS tumors and persist across time

For 51 of the 144 individuals in the study cohort, we collected biopsies of several non-contiguous tumors at study enrollment and/or biopsies of additional KS tumors at subsequent time points, most commonly 3 weeks (visit 2), 2 months (visit 5), and 6 months (visit 8) after study enrollment. We examined whether, in individual participants, T-cells from clustered unknown specificity groups that we hypothesize contain KSHV- or HIV-specific T-cells would be detected in non-contiguous tumors that were biopsied at the same time. We also examined whether, in individual participants, putative KSHV- or HIV-specific T-cells from clustered unknown specificity groups would be detected in KS tumors that were biopsied at different times during the 1-year period of observation after study enrollment. This analysis confirmed that TCR sequences belonging to clustered unknown antigenic specificity groups were consistently identified in non-contiguous KS tumors biopsied at the same time or at different times over the course of 1 year of observation. In both endemic (Figure 5A) and epidemic (Figure 5B) KS tumors, we observe remarkably stable populations of T-cells from clustered unknown specificity groups (blue and red bands respectively in Figure 3A, 3B) infiltrating distinct KS tumors sampled at various timepoints from study entry through the end of the one-year period of observation.

**Figure 5:**
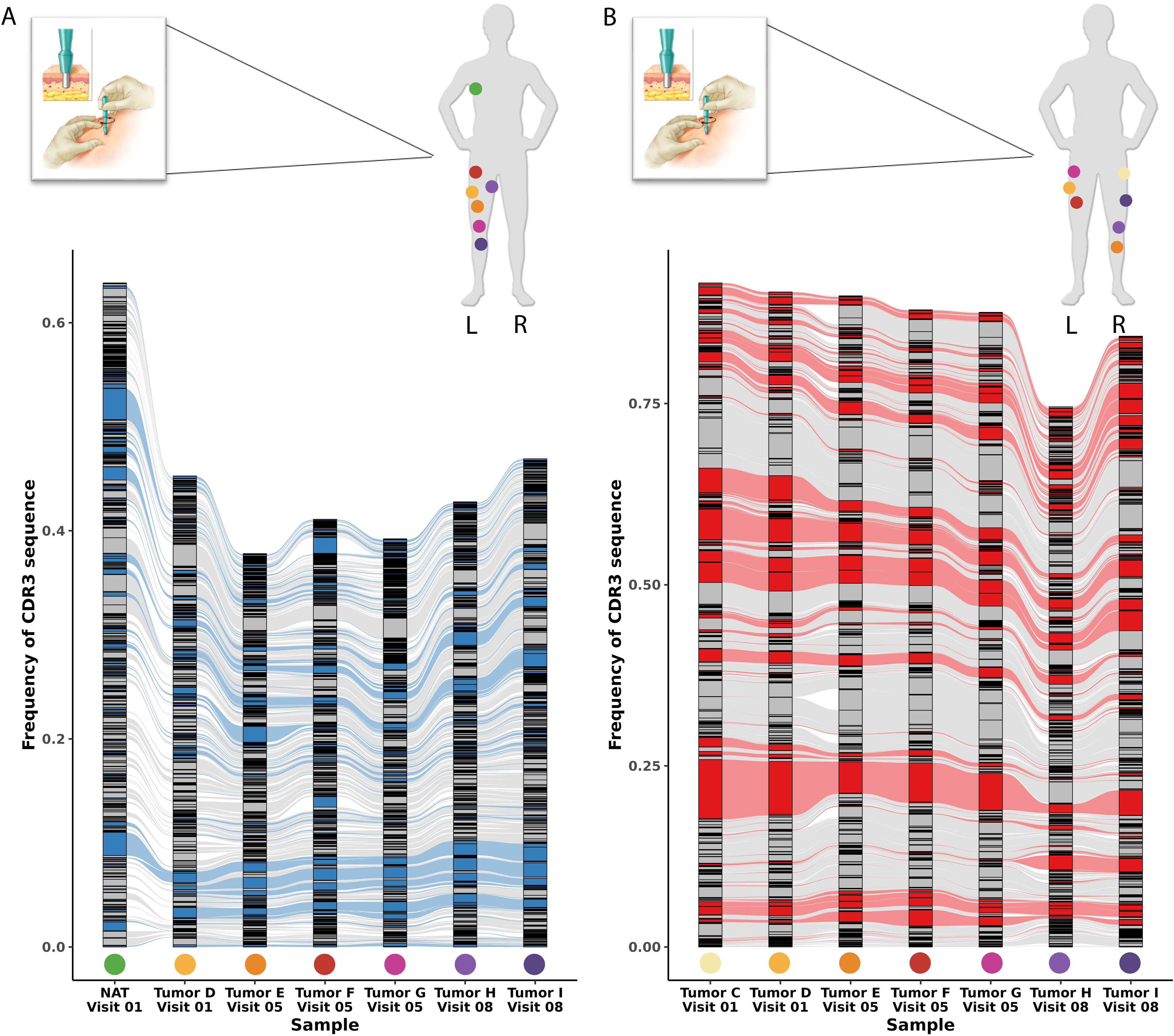
T-cells from GLIPH2-defined clusters with specificity for unknown antigens persist in the KS TME across time and space in individuals with epidemic and endemic KS. (A, B) Alluvial plots of the 500 most frequent T-cell receptor β chain sequences observed in tumor samples obtained from different locations on the body surface and at different time points in two representative individuals with (A) endemic KS and (B) epidemic KS. Highlighted are persistent TCR β chain CDR3 sequences from GLIPH2-defined “clustered unknown” antigenic specificity groups in endemic (blue) or epidemic (red) KS lesions. Colored dots on the body figures above each alluvial plot denote the sites from which each tissue sample was collected and are reproduced below the corresponding column containing the repertoire data for that tissue sample.

On average these T-cells were found at a cumulative frequency of 30%, 26% and 23% in tumors from individuals with endemic KS at visits 1, 5 and 8, respectively, and comprised 31%, 27% and 25% of the repertoire in tumors from individuals with epidemic KS collected at visits 1, 5, and 8, respectively.

### T-cells in clustered unknown specificity groups are antigen-experienced effector cells

The observation that clustered unknown T-cells carrying identical TCR β chain sequences were consistently observed in non-contiguous tumors suggested that they can circulate in the blood. Therefore, to define the phenotype of T-cells assigned to clustered unknown specificity groups and capture their paired TCR α and β chain sequences at large scale, we performed scRNA-seq with TCR α/β chain sequencing on 20 PBMC samples obtained at multiple time points after study entry from 9 individuals in our Uganda KS cohort (5 with epidemic KS, 4 with endemic KS). The datasets were normalized, integrated, and mapped to a reference PBMC single-cell atlas (13), creating a combined dataset comprising 246,521 single cells, of which 61,763 cells carry a productive αβ TCR (mean, 3,087 such cells per sample), collectively carrying 31,576 unique αβ TCRs. Of these 61,763 cells, 38,411 (62%) carrying 9,805 unique productive αβ TCRs could be presumptively mapped back to the KS TIL repertoire by virtue of their TCR β chain sequences, and 14,698 of those cells, carrying 4,283 unique αβ TCRs, belonged to “clustered unknown” antigenic specificity groups defined by GLIPH2 analysis.

Cell-type classification using Celltypist (14, 15) of the 61,763 cells carrying a productive αβ TCR revealed that 60,779 (98.40%) were classified as T-cells and 686 (1.1%) as NK cells (Figure 6A).

**Figure 6:**
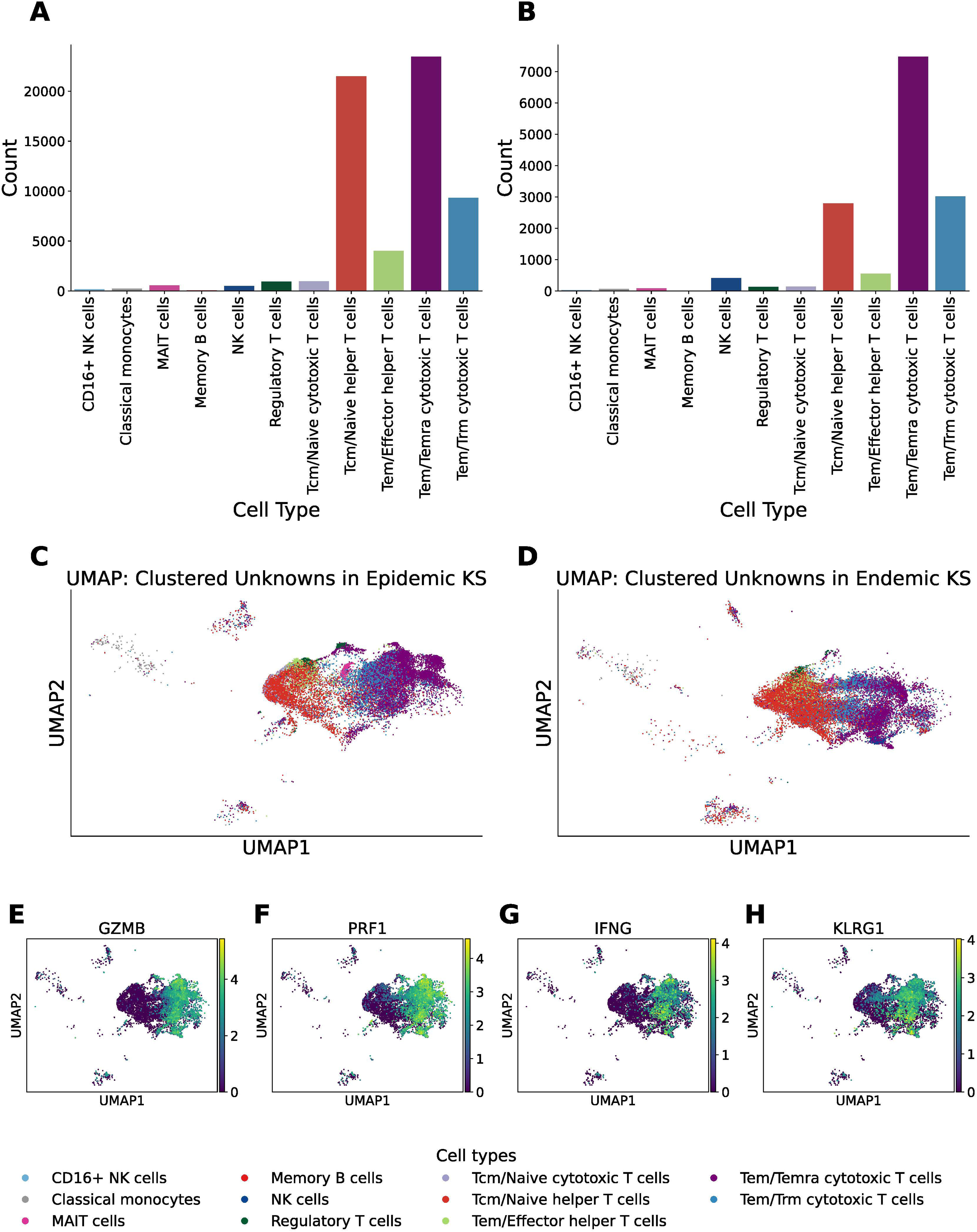
T-cells in “clustered unknown” antigenic specificity groups are primarily CD8^+^ effector memory cells. (A) Cell type classification using Celltypist (14, 15) of 61,763 cells carrying a productive αβ TCR from 20 single-cell gene expression and TCR VDJ libraries generated from PBMCs from 9 individuals with KS (4 endemic and 5 epidemic). (B) Cell type classification of the subset of 14,698 T-cells carrying a productive αβ TCR that were assigned to “clustered unknown” antigenic specificity groups by GLIPH2. (C, D) Integrated single-cell UMAP incorporating cell type classification of (C) 6,511 clustered unknown T-cells from the 5 individuals with epidemic KS and (D) 8,187 clustered unknown T-cells from the 4 individuals with endemic KS. (E-H) Gene expression profiles from the integrated single-cell UMAP of 14,698 T-cells carrying clustered unknown TCRs for *GZMB*, *PRF1*, *IFNG*, and *KLRG1*.

The remaining 289 cells were classified as other types of immune cells despite carrying a productive αβ TCR, which is likely due to inadequate coverage of marker genes used for classification. Similarly, 14,179 of the 14,698 (96.4%) cells carrying a productive αβ TCR that mapped back to the KS TIL repertoire and were assigned to a “clustered unknown” antigenic specificity group by GLIPH2 were classified as T-cells, with 440 (2.9%) classified as NK cells (Figure 6B). Further classification of the T-cells into subsets by helper or cytotoxic phenotype revealed that effector memory or effector memory/CD45RA T-cells with cytotoxic phenotypes were the most common T-cells in both the total population of 61,763 T-cells captured by scRNA-seq and the subset of 14,698 that mapped back to the KS TIL repertoire and were assigned to clustered unknown specificity groups. T-cells classified as cytotoxic phenotype comprised a larger proportion of the clustered unknowns in KS TIL (72.3%) than they did of the entire T-cell population (54.7%).

The transcriptional profiles of T-cells in PBMC from participants with epidemic KS that mapped back to KS tumors by virtue of their T-cell receptor β chain sequences were similar to those of corresponding T-cells from participants with endemic KS (Figure 6C, 6D). Differential gene expression analysis of T-cells carrying clustered unknown TCRs from participants with epidemic and endemic KS showed no significantly differentially expressed genes. T-cells classified as cytotoxic effector memory/CD45RA expressed high levels of granzyme B, perforin, and interferon-γ, but also high levels of *KLRG1*, which is associated with senescence and reduced proliferative capacity (Figure 6E-6H). Subsets of this group of T-cells expressed high levels of genes including *PDCD1*, *TIGIT*, *LAG3*, *HAVCR2*, and *TOX* that are associated with chronic stimulation and exhaustion (Supplementary Figure 3).

### Public TCRs from clustered unknown specificity groups recognize KSHV- and HIV-encoded peptides

To test the hypothesis that clustered unknown specificity groups are enriched for T-cells that recognize KSHV- or HIV-encoded peptides, we transduced T-cells with representative public clustered unknown TCRs from among the 4,283 unique TCRs that we captured by scRNA-seq and determined their antigenic specificity. We anticipated that TCRs classified as clustered unknowns that were only observed in samples from participants living with HIV infection would likely to be specific for antigens encoded by HIV, and that those observed in participants with or without HIV infection were possibly specific for antigens encoded by KSHV. To establish proof of principle for this approach, we first confirmed the predicted specificity and MHC restriction of an αβ TCR that our analytic pipeline had assigned to a clustered *known* specificity group. This αβ TCR carried a β chain encoding the CDR3 amino acid sequence CASSVDKGGTDTQYF, which has been associated with an HLA-B*42:01-restricted CD8**^+^** T-cell response to the HIV Pol_982-990_ peptide with sequence IIKDYGKQM (16). This TCR β chain was observed in multiple KS tumors from 13 individuals in our Uganda KS cohort (mean frequency, 0.35%), all of whom were HIV**^+^** and HLA-B*42:01**^+^**. Jurkat reporter T-cells (17, 18) and primary CD8**^+^** T-cells transduced with a αβ TCR carrying this β chain showed high-avidity recognition of HLA-B*42:01**^+^** target cells pulsed with Pol_982-990_ peptide (Figure 7A. 7B).

**Figure 7:**
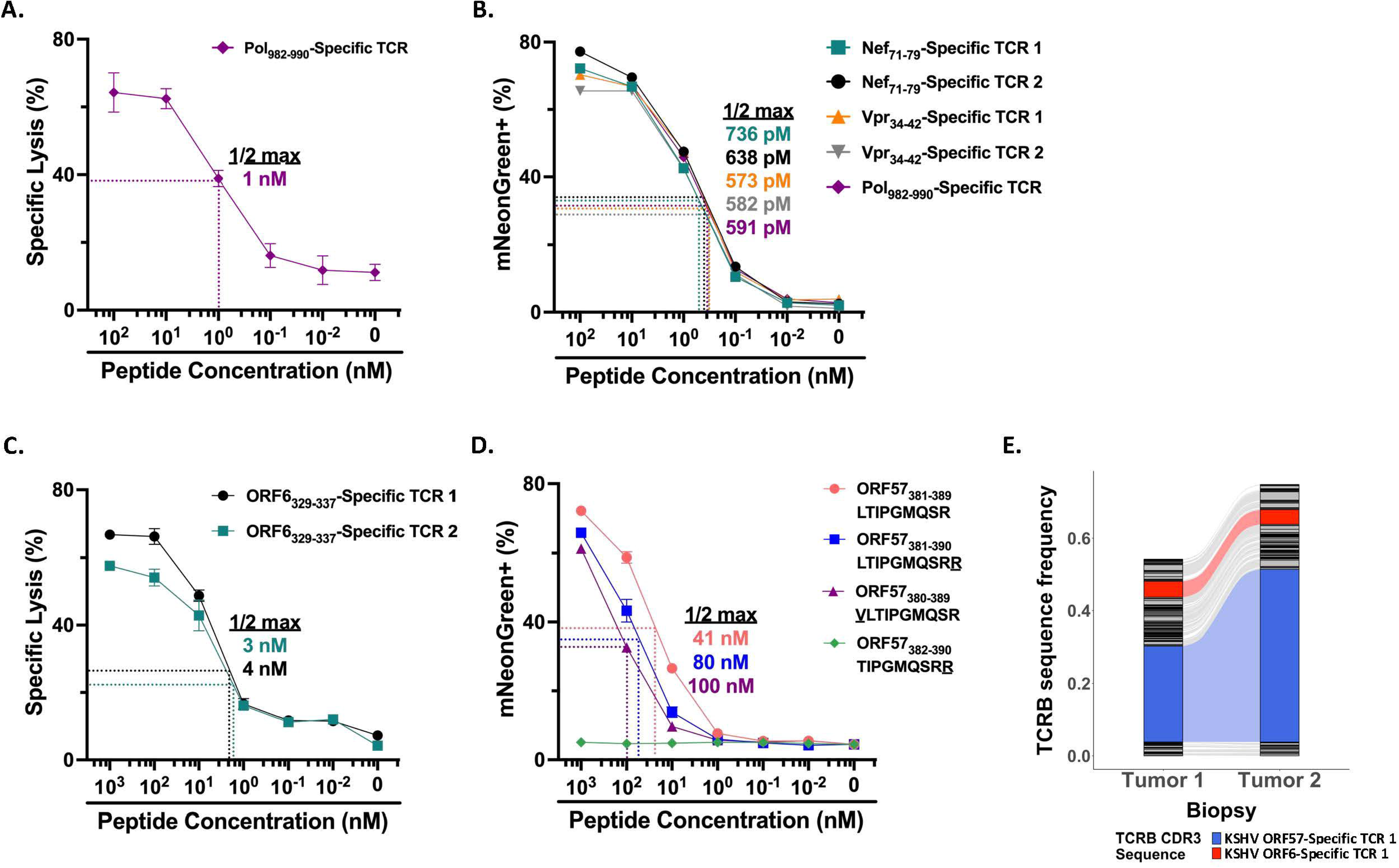
TCRs from clustered known and clustered unknown antigenic specificity groups recognize KSHV- and HIV-encoded antigens with high avidity. (A) Cytotoxicity of primary CD8**^+^** T-cells transduced with an αβ TCR specific for HIV Pol_982-990_/HLA-B*42:01 against HLA-B*42:01**^+^** EBV-transformed B-cells (EBV-LCL) pulsed with serial dilutions of HIV Pol_982-990_ peptide in a standard 4-hour ^51^Cr release assay. (B) Flow cytometric analysis of NeonGreen expression in Jurkat reporter T-cells transduced with αβ TCRs specific for HIV Nef_71-79_/HLA-B*42:01, HIV Vpr_34-42_/HLA-B*42:01, or HIV Pol_982-990_/HLA-B*42:01 after 16-hour co-culture with HLA-B*42:01**^+^**EBV-LCL pulsed with the corresponding peptides at the indicated concentrations. (C) Cytotoxicity of primary CD8**^+^** T-cells transduced with either of two related clustered unknown αβ TCRs against HLA-B*45:01**^+^**EBV-LCL pulsed with serial dilutions of KSHV ORF6_329-337_ peptide in a 4-hour ^51^Cr release assay. (D) Flow cytometric analysis of NeonGreen expression in Jurkat reporter T-cells transduced with a clustered unknown αβ TCR after 16-hour co-culture with HLA-A*66:01**^+^**EBV-LCL pulsed with serial dilutions of KSHV ORF57_381-389_, ORF57_381-390_, ORF57_380-389_, and ORF57_382-390_ peptides. (E) Alluvial plot of the 100 most frequent TCR β chain sequences in two non-contiguous KS tumors from HIV-seronegative HIPPOS participant 008_098. The red and blue-highlighted rivulets identify the TCR β chain sequences carried in the ORF6_329-337_- and ORF57_381-389_-specific TCRs featured in (C) and (D), respectively.

Similarly, an αβ TCR with β chain CDR3 amino acid sequence CASSLWAGGSNEQFF, which was also assigned to a clustered known specificity group and has been associated with an HLA-B*42:01-restricted CD8**^+^** T-cell response to the HIV Vpr_34-42_ peptide FPRPWLHGL (16), showed high-avidity recognition of HLA-B*42:01**^+^** target cells pulsed with Vpr_34-42_ peptide (Figure 7B, Vpr_34-42_-Specific TCR 1). In analogous studies we defined the antigenic specificity of four additional clustered unknown TCRs that we hypothesized to be HIV-specific. Three such TCRs from a single clustered unknown specificity group were found to recognize HIV Nef_71-79_ presented by HLA-B*42:01, and another putative HIV-specific TCR was found to recognize HIV Vpr_34-42_ presented by HLA-B*42:01 (Figure 7B).

To identify antigens recognized by putative KSHV-specific TCRs from clustered unknown specificity groups, we tested TCR-transduced T-cells for recognition of COS-7 cells that had been co-transfected with (i) pools of plasmids collectively encoding all annotated open reading frames (ORFs) in the KSHV genome, and (ii) a plasmid encoding the predicted MHC restricting allele. Plasmids encoding the KSHV ORFs in positive pools were individually tested for recognition to identify the ORF encoding the antigen. Iterative testing of successively smaller minigenes derived from that ORF was then performed to define the minimal antigenic peptide, with confirmation provided by testing recognition of synthetic peptide presented by target cells expressing the restricting class I MHC molecule. Two closely related clustered unknown TCRs from the same antigenic specificity group were found to recognize the KSHV ORF6_329-337_ peptide, AEQALHIGA, presented by HLA-B*45:01 (Figure 7C). These two TCRs were observed in multiple KS tumors from 14 participants in our Uganda KS cohort, all of whom were HLA-B*45:01**^+^**. Another clustered unknown TCR was found to recognize the KSHV ORF57_381-389_ peptide LTIPGMQSR presented by HLA-A*66:01, with slightly less avid recognition of the ORF57_381-390_ and ORF57_380-389_ peptides, but no recognition of the ORF57_382-390_ peptide (Figure 7D). This TCR comprised 23% and 43 %, respectively, of the TCRs in two KS tumors from one HLA-A*66:01**^+^**individual (Figure 7E).

## Discussion

Kaposi sarcoma most often develops years, and usually decades, after acquisition of primary KSHV infection in individuals with T-cell deficiency or dysfunction – such as people living with HIV or recipients of allogeneic hematopoietic or solid organ transplants. These observations suggest that loss or impairment of a T-cell component of pre-existing KSHV-specific immunity predispose to KS. Strategies that preserve or restore the T-cell component of KSHV-specific immunity in PLWH and others at risk should, therefore, have potential for the prevention or treatment of KSHV-associated disease.

We hypothesized that T-cells specific for KSHV, and possibly for HIV in PLWH who develop KS, would be attracted to KS tumors, and therefore performed comprehensive multi-omic analysis of the repertoire of T-cells found in 299 KS tumor biopsies from a cohort of 144 Ugandan adults presenting to the Uganda Cancer Institute. RNA-seq and targeted gene expression analysis demonstrated consistent expression of both latent and lytic KSHV genes in a subset of our KS tumors as well as in tumors studied by other groups (3, 4), and also detected consistent expression of HIV genes in the subset of tumors from PLWH. Surprisingly, RNA-seq analysis also suggested expression of a smaller, variable subset of HIV genes in 4 of 12 endemic KS tumors; we believe that this observation is an artefact of the *k*-mer-based strategy used for quantifying gene expression from the human and HIV transcriptomes.

The immune cell composition of the KS tumors is dominated by M2-polarized macrophages and CD8**^+^** T-cells. M2 macrophages were significantly more frequent in endemic than in epidemic tumors. We observed a negative correlation between the inferred content of M2 macrophages and CD8**^+^** T-cells, consistent with the possibility that M2 macrophages might somehow inhibit CD8**^+^** T-cell infiltration into or proliferation and persistence within the KS TME. Comprehensive AIRR-seq of our 299 KS tumors identified 2.85 million TCR β chain sequences that we believe capture a significant segment of the TCR repertoire of T-cells found in KS tumors in Ugandan adults. Less than 1% of these TCR β chain sequences are noted in existing TCR databases that catalogue the TCR variable region sequences that have been associated with T-cell responses to defined pathogen or tissue antigens. The diversity of the TCR repertoires of KS tumors is consistently higher in endemic than in epidemic tumors across a broad range of diversity metrics, which is likely due to the relative paucity of CD4**^+^** T-cells in epidemic tumors.

Our findings provide compelling support for the hypothesis that the KS tumors attract T-cells specific for KSHV and, in PLWH, HIV. Analysis with GLIPH2 of the 2.85 million TCR β chain sequences observed in the 299 KS tumors found that 24% of these sequences could be grouped into clusters of closely related TCR β chain sequences that are predicted to be (i) components of αβ TCRs with similar or identical, but unknown, antigenic specificity, and (ii) closely associated with a specific, known MHC allele. Within individual participants, T-cells carrying identical TCR β chain sequences and assigned to clustered unknown specificity groups were observed to associate with non-contiguous KS tumors, sometimes far apart on the body surface, that were biopsied either at the same time or at different times over the course of a year. Thus, these T-cells were capable of trafficking to non-contiguous tumors via circulation in the blood. We therefore performed scRNA-seq of PBMC from 9 of the 144 participants with KS and identified 14,698 T-cells carrying 4,283 unique αβ TCRs that could be confidently mapped back to KS tumors by virtue of their TCR β chain sequences. Transcriptional profiling of these 14,698 T-cells revealed that they were primarily antigen-experienced effector cells with a cytotoxic phenotype. Finally, we transduced T-cells with 9 different clustered unknown TCRs and showed that they recognized antigens encoded by KSHV ORF6 and ORF57 as well as HIV Nef and Vpr, presented by the predicted MHC allele for each TCR, with high avidity. The 4,283 unique αβ TCRs from clustered unknown specificity groups that we have identified likely contain many additional KSHV-specific and HIV-specific TCRs. Identification of the antigens recognized by additional putative KSHV-specific TCRs is the focus of current effort in our laboratory.

Our results provide a blueprint for the development of antigen-specific immune therapy for KS and possibly other KSHV-associated diseases. Whole exome sequencing by our group of 78 KS tumors and matched normal control skin from 59 adults with KS has recently revealed a very low mutational burden in all but one tumor specimen and no recurrent mutations (8). Transcriptional profiling of KS tumors by our group and others (2–4, 19), however, has revealed consistent expression of both latent and lytic cycle KSHV genes in KS tumors. Thus, our finding that KSHV-specific T-cells are attracted to KS tumors offers the prospect of antigen-specific therapy for KS based on vaccination or adoptive T-cell transfer. Indeed, adoptive therapy with either autologous or allogeneic EBV-specific T-cells has been used for >25 years to treat EBV-associated lymphoma (20), and tabelecleucel, an off-the-shelf allogeneic EBV-specific T-cell immunotherapy (21, 22), has become standard of care in the European Union for the treatment of EBV**^+^** posttransplant lymphoproliferative disease.

Developing effective therapy for KS that is based on recognition and elimination of KS tumor cells by KSHV-specific T-cells will require answering several critically important questions. Can CD8**^+^** KSHV-specific T-cells recognize and eliminate KSHV-infected cells? Which of the KSHV-encoded antigens previously identified and studied by other groups (23–36) and ours are the optimal target antigens? Are there cells in KS tumors that are not infected with KSHV that will nonetheless require elimination for complete tumor eradication? Will KSHV-specific T-cells elicited by therapeutic vaccination or infused after *ex vivo* manipulation be able to penetrate into KS tumors adequately? What are the mechanisms and maneuvers that KSHV exploits to evade recognition and elimination by T-cells, and how can these be thwarted or counteracted? For example, is the prominent population of M2 macrophages in KS tumors induced in some way by KSHV (37), and do these macrophages inhibit the proliferation or function of KSHV-specific T-cells, or impair or prevent their entry into the KS TME (38)? Does HIV in KS tumors induce or potentiate the M2 polarization of intratumoral macrophages? Answering these questions is a high priority for the field.

Further studies of the KSHV-specific T-cell response offer the prospect of developing immune interventions for preventing KS and other KSHV-associated diseases, particularly in PLWH who are at greatest risk. The strong association of KS with T-cell deficiency or dysfunction suggests that this T-cell response is impaired or disabled in individuals who develop the disease. Comparison of the properties of KSHV-specific T-cells from asymptomatic KSHV-seropositive individuals, including children with recently established immunity and adults for whom primary KSHV infection is a remote event, and those from individuals with KS could potentially enable identification of those in whom pre-emptive interventions designed to restore or enhance KSHV-immunity could forestall development of clinical disease.

## Materials and Methods

### Subjects and samples

Adults 18 years or older with histologically confirmed Kaposi sarcoma who were naïve to ART and had received no prior chemotherapy were enrolled between October 2012 and May 2021 on a prospective cohort study (termed “HIPPOS”) at the Uganda Cancer Institute (UCI) in Kampala, Uganda, as previously described (39). Participants completed an enrollment visit and up to 10 follow-up visits over ∼1 year. At enrollment, participants completed a standardized medical history and physical examination, a blood sample and an oral swab were collected for KSHV testing, and peripheral blood mononuclear cells (PBMC) were isolated. HIV serology was performed, and CD4**^+^** T-cell and plasma HIV RNA testing was completed for HIV-seropositive participants. UCI medical charts were reviewed to record routine blood count data and chest X-ray findings. Full-thickness punch biopsies (4 x 4 mm diameter) of at least three non-contiguous cutaneous KS lesions and a punch biopsy of axillary skin not visibly involved with KS were also obtained and placed in either neutral buffered formalin or RNA*later* (Thermo Fisher Scientific, Waltham, MA, USA).

At follow-up visits, participants completed an interim medical history and detailed physical exam to assess treatment response and provided additional blood samples and oral swabs for KSHV testing. Up to three additional biopsies of non-contiguous KS lesions and a blood sample for PBMC isolation were obtained at follow-up visits at 3-, 36-, and 48-weeks following enrollment, and placed in RNA*later*.

Participants were offered chemotherapy at the UCI per standard guidelines independent of the study. First-line therapy consisted of combination bleomycin and vincristine (BV) or paclitaxel; both regimens were given every 3 weeks for six cycles. HIV-seropositive participants not already in HIV care were referred to local HIV clinics for management and initiation of ART according to Ugandan Ministry of Health guidelines.

Biopsies of Burkitt lymphoma (BL) were acquired from 2 cohorts of African children after obtaining verbal or written informed consent using institutional review board–approved protocols, as previously described (7). The first cohort comprised 12 children with BL who presented to the Uganda Cancer Institute (UCI) in Kampala, Uganda between August 2013 and March 2014. The second cohort comprised 37 children diagnosed with BL between 1975 and 1992 at Korle Bu Hospital in Accra, Ghana, from whom cryopreserved, archival BL tumor biopsies were available.

### Laboratory procedures for viral and T-cell monitoring

Plasma and oral swab samples were evaluated for KSHV DNA by quantitative, real-time PCR at the Hutchinson Centre Research Institute – Uganda Laboratory in Kampala, Uganda as previously described (2, 39). Samples with >150 copies/mL of KSHV DNA were considered positive (40). CD4**^+^** T-cell counts were measured by flow cytometry, and HIV-1 RNA levels were measured using real-time RT-PCR.

### RNA sequencing (RNA-seq)

Total RNA was extracted from cryopreserved KS tumor biopsies using the RNeasy Fibrous Tissue Mini Kit (QIAGEN, Venlo, The Netherlands), quantitated on a Qubit 3.0 Fluorometer (Thermo Fisher), and assessed on an Agilent 2200 Bioanalyzer (Agilent Technologies, Inc., Santa Clara, CA). RNA samples with RNA integrity number (RIN) greater than 7 were used for sequencing library preparation. Sequencing libraries from RNA extracted from cryopreserved KS tumor biopsies from 39 HIV-seropositive HIPPOS participants (epidemic KS) were prepared in the Fred Hutchinson Cancer Center (FHCC) Genomics Core Facility (Seattle, WA) using the TruSeq RNA Sample Prep Kit v2 (Illumina, San Diego, CA, USA), as previously described (2). Library size distributions were validated using the Agilent 2200 TapeStation. Library QC, pooling of indexed libraries, and cluster optimization were performed using the Qubit 3.0 Fluorometer. Pooled libraries were sequenced on an Illumina HiSeq 2500 in “High Output” mode with a paired-end, 50-base read configuration. Sequencing libraries from RNA extracted from cryopreserved KS tumor biopsies from an additional 12 HIV-seronegative HIPPOS participants were prepared and sequenced at Azenta (South Plainfield, NJ) using the NEBNext Ultra II RNA Library Prep Kit for Illumina (New England Biolabs, Ipswich, MA, USA). The sequencing libraries were validated on the Agilent TapeStation, quantified using the Qubit 3.0 Fluorometer as well as by quantitative PCR (KAPA Biosystems, Wilmington, MA, USA), and sequenced on a partial S4 lane of an Illumina NovaSeq 6000 with a paired-end, 150-base read configuration.

### Targeted gene expression analysis

RNA was extracted from formalin-fixed paraffin-embedded (FFPE) slides from 25 epidemic KS and 2 endemic KS tumor biopsies and 5 samples of uninvolved skin using the AllPrep DNA/RNA FFPE Kit (QIAGEN). Gene expression analysis was performed on RNA samples on the Nanostring platform (Nanostring Technologies, Seattle, WA, USA) using the Immunology V2 Panel supplemented with 30 user-defined viral and host gene probes. Of the 25 epidemic KS tumors profiled, 8 were also profiled using bulk RNA-seq. Raw probe counts were normalized using the geometric mean of 15 housekeeping genes followed by analysis using nSolver software (Version 4.0) and the nSolver Advanced Analysis package (version 2.0.115).

### T-cell receptor β chain variable region sequencing

Genomic DNA was extracted from 363 cryopreserved biopsies of KS tumor and normal skin (termed normal adjacent tissue, NAT) using the DNeasy Blood & Tissue Kit (QIAGEN) and prepared for T-cell receptor β chain (*TRB*) variable region sequencing at survey level resolution (41) using the ImmunoSEQ hsTCRB v3.0 assay (Adaptive Biotechnologies, Seattle, WA). *TRB* sequencing was performed on pools of libraries from 11 - 15 tumor or skin samples on the Illumina MiSeq platform using a v3 150-cycle kit (Illumina, San Diego, CA) in the FHCC Genomics Core Facility (Seattle, WA) or the Hutchinson Centre Research Institute – Uganda Laboratory (Kampala, Uganda).

### MHC genotyping

High-resolution genotyping of the MHC class I and class II loci of all participants in the epidemic and endemic KS cohorts and the Uganda BL cohort was performed on genomic DNA via next-generation sequencing as a contract service by Scisco Genetics, Seattle, WA. The full standardized protocol is available online (42). Sequencing of the PCR amplicons spanning the MHC loci was performed on the Illumina MiSeq platform using a v2 500-cycle kit or a v2 500 nano-cycle kit (Illumina, San Diego, CA) in the FHCC Genomics Core Facility or the Hutchinson Centre Research Institute – Uganda Laboratory.

### Single-cell RNA sequencing (scRNA-seq)

Viable cells were thawed from cryopreserved PBMC specimens and subjected to scRNA-seq using the 10X Genomics (10X Genomics, Pleasanton, CA) platform. Single index chemistry was performed for all libraries using the following reagents to generate scRNA-seq libraries for V1.0 chemistry: the Chromium^TM^ Single Cell 5’ Library Construction Kit, Chromium ^TM^ Single Cell A Chip Kit, ChromiumTM Single Cell 5’ Library & Gel Bead Kit, Chromium^TM^ Single Cell 5’ Feature Barcode Library Kit, and the Chromium^TM^ Single-Cell Enrichment, Human T-Cell kit. For libraries generated using V1.1 chemistry, the Chromium^TM^ Next GEM Single Cell 5’ Library and Gel Bead Kit v1.1, and the Chromium^TM^ Next GEM Chip G Single Cell Kit were used.

Libraries were quantified using the Qubit 3.0 Fluorometer (Thermo Fisher, Waltham, MA), TapeStation using the Agilent TapeStation (Agilent, Santa Clara, CA), and by qPCR using the KAPA Library Quantification Kit for Illumina (Roche, Basel, Switzerland). Libraries were pooled and sequenced on the Illumina NovaSeq 6000 (Illumina, San Diego, CA) on an SP1 flow cell in the FHCC Genomics Core.

### Analysis of RNA-seq data

A total of 4,353,735,624 paired-end reads were obtained from 51 RNA-seq datasets (Mean: 85,367,365 reads per sample) generated from 39 epidemic KS and 12 endemic KS tumor samples. We used BBDuk (43) to remove any adapter contamination using a starting k-mer size of 23 and min-k size of 11, with an allowed Hamming distance of 1 between query k-mer and reference adapter k-mers. K-mers with average quality less than 6 were trimmed from both ends of the read. Reads with length ≤ 10bp were discarded after trimming. We quantified the transcript and gene level expression against the human (Gencode v42), HHV8 (GCF_000838265.1), and HIV (GCF_000864765.1) transcriptome index using Salmon v1.9.0 (44). K-mer sizes of 31 for the human transcriptome and 21 for the viral transcriptomes (HIV and HHV8) were used to build the reference index and quantify gene expression from the filtered reads.

We generated a gene-expression matrix from the Salmon gene-level expression data with sample name as rows, gene ensemble ID as columns, and expression values in each cell represented in transcripts per million reads (TPM). We quantified the cell-type composition of the tumor samples using the LM-22 immune cell-type model from CIBERSORTx (5, 45).

We included in our analysis publicly available RNA-seq data generated from 6 paired epidemic KS tumor –Normal Adjacent Skin (NAT) samples (3) and 18 paired epidemic KS tumor - NAT samples, 6 paired endemic KS tumor - NAT samples, and 3 control skin samples (4) to validate the findings from our cohort. The raw dataset was downloaded from NCBI SRA using the NCBI SRA-Toolkit. The 59 single-end RNA-seq datasets from the two sources were processed using the same BBMAP-Salmon pipeline modified to accommodate single-end sequencing datasets. Gene expression matrices with TPMs for 30 sun-exposed and 30 non-sun-exposed control skin samples were obtained from the from the Genotype-Tissue Expression (GTEx) project (6), and used to assess the baseline transcriptomic profile of normal skin. The GTEx data were accessed through the GTEx Portal on April 27, 2023. These 119 RNA-seq quantification datasets from 6 sequencing projects were formatted for analysis with CIBERSORTx (5, 45), and the immune cell-type composition for each dataset was determined using the LM-22 immune cell-type model.

### T-cell receptor repertoire analysis

T-cell repertoire analyses were conducted using the R statistical programming language. A reproducible workflow of all steps in data QC, data transformation, and analysis was developed using Jupyter notebooks and the Nextflow workflow management system (46). Repertoire analysis was performed with open-source R packages, and custom scripts were implemented in LymphoSeq2, an R package that we developed for the exploration of Adaptive Immune Receptor Repertoire Sequencing (AIRR-Seq) data (47).

To investigate intratumoral T-cell heterogeneity, we sequenced a total of 474 TCR repertoires from both tumor and adjacent normal tissues collected from 146 individuals enrolled in the study. Repertoires with less than 1,000 total *TRB* sequences were removed from the analysis, leaving a total of 363 TCR repertoires from 299 KS tumor biopsies (221 epidemic KS, 78 endemic KS), 64 samples of uninvolved axillary skin (39 epidemic KS, 25 endemic KS), and 7 samples from single-cell tumor suspensions subjected to a Rapid Expansion Protocol (6 epidemic KS and 1 endemic KS). In addition, we included in the repertoire analyses 49 AIRR-Seq datasets generated from BL tumors from children from Uganda and Ghana using the same wet- and dry-lab platforms as the KS tumor and NAT tissues, which also passed the same sequencing threshold of 1,000 total *TRB* sequences.

To minimize the impact of differences in the effective depth of sequencing achieved on the samples on the assessment of repertoire diversity, repertoires were sampled down to a total count of 1,000 *TRB* nucleotide sequences (48). The sampled repertoires were aggregated over the unique *TRB* CDR3 nucleotide sequences and repertoire diversity statistics were averaged over 100 iterations.

Repertoire diversity was measured using Rényi generalized entropy at different values of α (α = 0, 0.25, 0.5, 1, 2, 4, 8, 16, 32, 64, ∞) to capture different characteristics of the T-cell repertoire diversity. Species richness (α = 0) and Shannon entropy (α = 1) are entropy measures that are more affected by the presence of distinct T-cells in the repertoire at any degree of abundance. Simpson’s diversity (α = 2) and Inverse Berger-Parker index (α = ∞) are entropy measures that weigh the more abundant T-cells in the repertoire to a greater extent. Hence, they give a summary of the diversity among expanded T-cells in the repertoire. We used a custom R wrapper around the Rényi function provided in the Vegan R package to calculate the depth-normalized Rényi diversity profile for each dataset.

### Prediction of T-cell antigenic specificity and MHC restriction

The TRB CDR3 amino acid sequences in each repertoire were compared with the VDJdb (9, 10, 49) and McPAS-TCR (11) databases of annotated TRB sequences, with high confidence of antigenic specificity (VDJdb score >1; McPAS-TCR antigen identification method < 3), to identify sequences that have previously been associated with a T-cell response to a specific microbial or tissue antigen or with a specific pathological condition. TCRs from public databases were filtered to ensure that the MHC allele associated with each TCR in the database was also carried in the individual with the TCR sequence of interest.

Antigenic specificity groups of T-cell receptors carrying specific TRB CDR3 amino acid sequences were computationally predicted using GLIPH2 (12). Sequences from T-cell receptor repertoires were labeled with repertoire ID and phenotype (epidemic KS, endemic KS). MHC typing information from participants in the HIPPOS cohort was formatted as per GLIPH2 requirements. A reference database for GLIPH2 was generated using 35 control AIRR-Seq PBMC datasets from sub-Saharan Africa. The custom reference catalogued the CDR3 length distribution, V-gene usage, and CDR3 composition of 932,116 unique TRB CDR3 sequences.

TRB CDR3 amino acid sequences with likely antigenic specificity for CMV, EBV, HIV, *Mycobacterium tuberculosis* (*Mtb*), or an identified tissue antigen, were identified by first finding all specificity groups that contained an annotated TRB sequence reported in one or more of the public databases. All TRB CDR3 amino acid sequences belonging to these specificity groups were identified as sequences with likely antigenic specificity for CMV, EBV, HIV, *Mtb*, a specific tissue antigen, or a specific pathological condition (48). GLIPH2 groups that did not contain any TRB CDR3 amino acid sequences with associated antigenic specificity were categorized as “Clustered Unknowns”. Sequences not grouped into antigenic specificity groups by GLIPH2 and with yet undetermined antigenic specificity were categorized as “Unclustered Unknowns”.

### Analysis of single-cell RNA-seq gene expression (GEX) and VDJ libraries

We built a structured analysis pipeline using NextFlow to process scRNA-seq GEX + VDJ sequencing libraries from PBMC samples of 20 individuals with epidemic and endemic KS. We quantified the number of cells across both GEX and VDJ libraries using the Cell Ranger multi (pipeline version: Cell Ranger-7.0.1). The references for Cell Ranger GEX (Human reference (GRCh38) - 2020-A) and VDJ (Human V(D)J reference (GRCh38) ensembl-7.0.0) were obtained from the 10X Cell Ranger website (https://www.10xgenomics.com/support/software/cell-ranger/downloads). For quality control and preprocessing, we utilized the Bioconductor package scuttle (50) to compute per-cell metrics. We used the isOutlier function from the Bioconductor package Scater (50) to identify outliers based on per-cell metrics, with a threshold of 5 median average deviations. Cell counts were normalized, and clustering was performed using scuttle and scran. Doublet clusters were identified based on gene expression using the scDblFinder (51). Seuratv4 (13), and scRepertoire (52) were employed to merge gene expression and VDJ data for each sample. T-cell sequence data from VDJ libraries were incorporated into the metadata field of the Seurat objects.

Gene expression and VDJ data from the 20 PBMC single-cell datasets were integrated using the Seurat rpca protocol (https://satijalab.org/seurat/articles/integration_rpca.html). Ensembl IDs were replaced with gene symbols to prepare the integrated object for cell type classification.

The subset of cells with productive VDJ rearrangements was identified, and their phenotypic characterization was performed using CellTypist (14, 15) with the “Immune All Low” model. This model defines the phenotypic characteristics of 98 immune sub-populations derived from 20 tissues across 18 studies.

The integrated Seurat object was subsete and stored as h5ad files using SeuratDisk. These files were loaded into the AnnData format using Scanpy (53), which is compatible with CellTypist. The raw counts in the AnnData object were log-normalized, and CellTypist annotation was conducted with majority voting enabled. Finally, UMAP coordinates were calculated using the Scanpy UMAP function.

### Antigen grouping and phenotypic characterization of scRNA-seq GEX and VDJ libraries

We used the SeuratV4 reference mapping protocol (https://satijalab.org/seurat/articles/multimodal_reference_mapping#intro-seurat-v4-reference-mapping) for cell-type classification of cells in the single-cell libraries utilizing a multimodal PBMC atlas comprising 161,764 cells as a reference (29). To ensure consistency with the reference atlas normalization, we first normalized cell counts within the integrated Seurat object using the scTransform function from Seurat. We identified anchors between the reference and query through the FindTransferAnchors function, transferring cell type labels using the MapQuery function. We merged GLIPH2 results from bulk TRB sequencing data with TRB sequences from single-cell datasets to identify antigen groups. The merged table was filtered to include only TRB sequences classified as “Clustered Unknowns” from the previous section.

### Identification of KSHV- and HIV-encoded antigens recognized by CD8^+^ T-cells

Lentiviral transfer plasmids encoding the complete α and β chain sequences of clustered unknown, putative KSHV- or HIV-specific TCRs, separated by a self-cleaving porcine teschovirus-1 2A sequence (P2A) sequence and downstream of the MSCV U3 promoter, in the pRRLSIN.cPPT.MSCV/WPRE lentiviral plasmid backbone, were synthesized by Azenta Life Sciences (South Plainfield, NJ). Recombinant lentiviruses encoding TCR αβ transgenes were manufactured from the transfer plasmids by the Vector Production Facility at the Fred Hutchinson Cancer Center and used to transduce Jurkat clone E6.1 cells carrying an NR4A1_NeonGreen fusion transgene and inactivated endogenous *TRA*/*TRB* genes (17, 18) or primary CD8**^+^** T-cells from healthy donors. Primary CD8^+^ T cells were transduced after activation with CD3/CD28 Dynabeads (Thermo Fisher Scientific) at a 3:1 ratio overnight in 1 mL CTL media (RPMI-1640 supplemented with 10% heat inactivated human serum, L-glutamine, 1% penicillin/streptomycin, and 50 µM β-mercaptoethanol) with 50 U/mL interleukin-2 (IL-2). T-cells were transduced with 1 µL lentiviral supernatant in the presence of 4.5 µg/mL polybrene (Millipore Sigma) by centrifugation at 800 x g for 90 minutes at 32 C followed by a half-media change after 8 hours. TCR-transduced Jurkat cells were used for antigen identification studies on day +4 after transduction or later, and primary CD8**^+^** T-cells were expanded in CTL media supplemented with IL-2 and used for antigen identification on day +9 after transduction or later.

A cDNA library of all annotated protein-coding ORFs (93 total) in the KSHV genome was assembled in the pDEST103 plasmid backbone (Invitrogen). ORFs used for assembly of this KSHV ORF library were purchased from Azenta (South Plainfield, NJ) or received as generous gifts from Dr. J. Haas (54) or Dr. M. Stürzl (55). KSHV ORF minigene plasmids were generated by cloning PCR-amplified segments from an ORF or directly annealed complementary oligonucleotides designed with 3’ overhanging deoxyadenosines, into pcDNA3.1/V5-His-TOPO (Invitrogen). The mCherry-2xCL-YFP-Vpr plasmid encoding HIV-1 Vpr in the pcDNA3.1/Zeo(+) backbone was obtained from Addgene (#105215). The pcDNA3.1 SF2 Nef plasmid encoding HIV-1 Nef was obtained from the NIH-AIDS Reagent Program (#11431).

Specificity of clustered unknown TCRs for antigens encoded by KSHV or HIV was established by testing Jurkat reporter T-cells transduced with clustered unknown αβ TCRs with COS-7 cells transiently co-transfected with (i) pools of 10 plasmids encoding 10 KSHV ORFs or with single plasmids encoding HIV ORFs, and (ii) a plasmid encoding the GLIPH2-predicted class I MHC restricting allele for each TCR. Transfection was conducted in 96-well flat-bottom plates seeded the previous day with 9 x 10^3^ COS-7 cells, using 150 ng/well of each KSHV or HIV ORF plasmid and 4.5 µL/well FuGENE 6 (Promega). TCR-transduced Jurkat reporters (9 x 10^3^/well) carrying the NR4A1_NeonGreen transgene and suspended in 200 µL CTL media were added 24 h after transfection. After 18 hours of co-culture with COS-7 cells, the Jurkat reporter T-cells were collected from each well and assessed for NeonGreen expression by flow cytometry. The KSHV ORFs from pools that stimulated the Jurkat reporters were tested individually to identify the single ORF encoding the antigen. Iterative testing of successively smaller minigenes derived from that ORF then defined the minimal antigenic peptide, with confirmation provided by testing recognition of synthetic peptide presented by EBV-transformed B-cells expressing the restricting class I MHC molecule. Cytotoxicity of TCR-transduced primary CD8**^+^** T-cells was assessed in standard 4-hour ^51^Cr release assays.

The amino acid sequences of the TCR α and β chains carried by the 9 KSHV- or HIV-specific TCRs that were studied, along with their inferred peptide specificity and class I MHC restriction, are detailed in Supplementary Table 1.

**Supplementary table 1.**

| TCR ID | TRB CDR3 Sequence | TRA CDR3 Sequence | MHC Restriction | Peptide |
| --- | --- | --- | --- | --- |
| **Pol<sub>982-990</sub>-Specific TCR** | **CASSVDKGGTDTQYF** | **CAVSGGGFQKLVF** | **HLA-B\*42:01** | **HIV-1 Pol<sub>982-990</sub>** |
| **Nef<sub>71-79</sub>-Specific TCR 1** | **CASSQEFPGALYNEQFF** | **CAHGSSNTGKLIF** | **HLA-B\*42:01** | **HIV-1 Nef<sub>71-79</sub>** |
| **Nef<sub>71-79</sub>-Specific TCR 2** | **CASSQEFPGAAYNEQFF** | **CVPLSNTGKLIF** | **HLA-B\*42:01** | **HIV-1 Nef<sub>71-79</sub>** |
| **Nef<sub>71-79</sub>-Specific TCR 3** | **CASSQEFPGASYNEQFF** | **CGTESSGGTSYGKLTF** | **HLA-B\*42:01** | **HIV-1 Nef<sub>71-79</sub>** |
| **Vpr<sub>34-42</sub>-Specific TCR 1** | **CASSLWGGPSNEQF** | **CAYSGAGSYQLTF** | **HLA-B\*42:01** | **HIV-1 Vpr<sub>34-42</sub>** |
| **Vpr<sub>34-42</sub>-Specific TCR 2** | **CASSLWAGGSNEQFF** | **CAPDRAGGTSYGKLTF** | **HLA-B\*42:01** | **HIV-1 Vpr<sub>34-42</sub>** |
| **ORF6<sub>329-337</sub>-Specific TCR 1** | **CASSIAGHEQYF** | **CAVGASGGYQKVTF** | **HLA-B\*45:01** | **KSHV ORF6<sub>329-337</sub>** |
| **ORF6<sub>329-337</sub>-Specific TCR 2** | **CASSIAGHEQFF** | **CAVAASGGYQKVTF** | **HLA-B\*45:01** | **KSHV ORF6<sub>329-337</sub>** |
| **ORF57-Specific TCR 1** | **CASSTGVYGYTF** | **CAAITGGGADGLTF** | **HLA-A\*66:01** | **KSHV ORF57<sub>381-389</sub>** |

### Statistical methods

The statistical significance of repertoire diversity differences between epidemic and endemic KS participants and BL participants from Ghana and Uganda was assessed the Wilcoxon rank-sum test for continuous variables. We used a generalized linear model (GLM) to examine the relationship between percentage abundance of CD8**^+^** T-cell and M2 Macrophages from RNA-seq datasets on KS tumors. The correlation between percent abundance of the two cell types in KS tumors was measured using a two-sided Pearson test.

### Ethical considerations

Approval for the study was obtained from the Makerere University School of Medicine Research and Ethics Committee, Fred Hutchinson Cancer Center IRB, and Uganda National Council for Science and Technology. All participants provided documentation of informed consent.

## Acknowledgements

We would like to acknowledge the HIPPOS study team, the Data and Laboratory Teams at the Hutchinson Center Research Institute – Uganda, HIPPOS study participants, and their families for enabling this research. We would also like to thank Dr. Jeffrey Vieira, Sydney Favors, Denise Tong, and Dr. Kerry J. Laing for assistance with generation and maintenance of the KSHV ORF plasmid library. Finally, we would like to thank The Genotype-Tissue Expression (GTEx) Project for data used in this manuscript: the Sun-exposed skin and Non-sun-exposed skin datasets from the v8 data release. The Genotype-Tissue Expression (GTEx) Project is supported by the Common Fund of the Office of the Director of the National Institutes of Health, and by NCI, NHGRI, NHLBI, NIDA, NIMH, and NINDS.

## Data Availability

AIRR-Seq datasets from the 363 KS tumor and NAT samples and 49 BL tumor samples from Ghana and Uganda are publicly available for access on the Adaptive Biotechnologies immunoSEQ Analyzer portal at (https://clients.adaptivebiotech.com/pub/ravishankar-2024-jem) https://clients.adaptivebiotech.com/pub/. All R scripts, metadata, GLIPH2 references and Nextflow workflows to analyze and visualize the data used in the study are available on GitHub (https://github.com/shashidhar22/ks_manuscript). Data generated by Nextflow pipelines and GLIPH2 used to generate figures for the manuscript are available on Zenodo (https://zenodo.org/records/10594758). RNA-seq datasets from 51 KS tumors and scRNA-seq GEX+VDJ datasets from 20 KS PBMC libraries are available through the NCBI under BioProject number PRJNA1068629 (https://www.ncbi.nlm.nih.gov/bioproject/PRJNA1068629).

## Ethics Statement

The studies involving human participants were reviewed and approved by the Makerere University School of Medicine Research and Ethics Committee, the Fred Hutchinson Cancer Center IRB, and the Uganda National Council for Science and Technology. All participants provided documentation of informed consent.

## Author Contributions

EW, WP, AT, and SR conceived and designed the study. PM, JK, JN, SS, and DM collected the tissue samples. AT, LO, PA, and JO generated the AIRR-Seq and MHC genotyping data. LA generated and analyzed the targeted gene expression data. JW, DK, and LJ contributed to the KSHV ORF plasmid collection. IT completed the T-cell antigen identification studies, with assistance from CM. SR, DC, AT, and EW analyzed the data. SR, AT, IT, and EW drafted the manuscript. All authors edited the manuscript. All authors contributed to the article review and approved the submitted version.

## Funding

This research was supported by awards from the National Cancer Institute (R01 CA217138, R01 CA239287, P30 CA015704-043S3) and from the Immunotherapy and Pathogen-Associated Malignancies Integrated Research Centers at the Fred Hutchinson Cancer Center. This research was also supported by the Specimen Processing and Research Cell Bank and the Genomics and Bioinformatics Shared Resources of the Fred Hutch/University of Washington Cancer Consortium (P30 CA015704). Bioinformatic analysis was supported by the Scientific Computing Infrastructure at the Fred Hutchinson Cancer Center funded by ORIP grant S10OD028685.

## Conflict of Interest

The authors declare that the research was conducted in the absence of any commercial or financial relationships that could create a conflict of interest.

## Notes

https://github.com/shashidhar22/ks_manuscript

### Competing Interest Statement

The authors have declared no competing interest.

### Summary of Updates

Updated author list and affiliations; Author contributions updated;

