## Supplementary figures and images for "T-cells specific for KSHV and HIV migrate to Kaposi sarcoma tumors and persist over time"

### Supplementary Figure 1

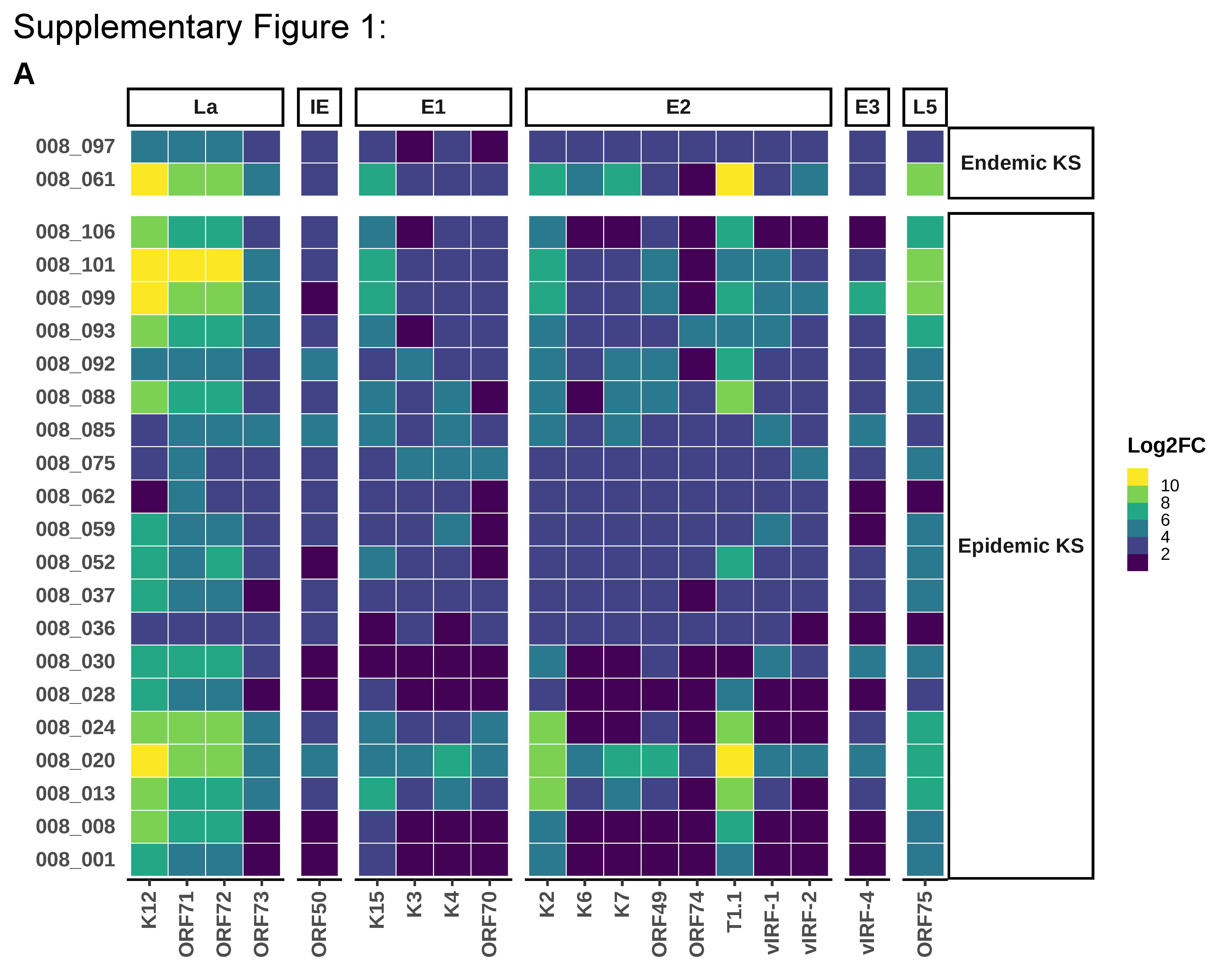

### Supplementary Figure 2

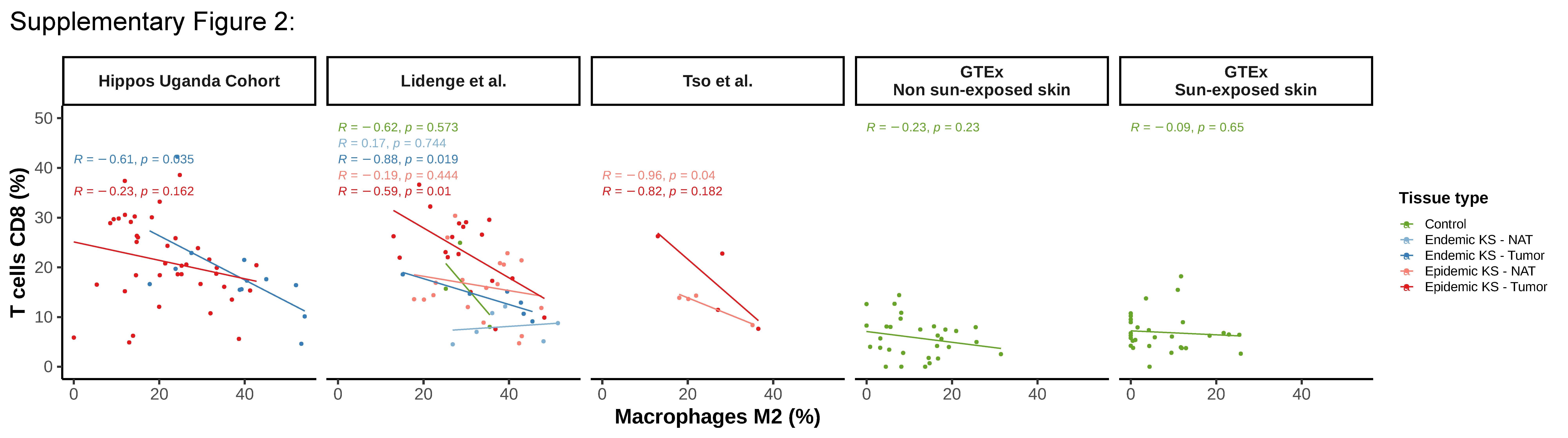

### Supplementary Figure 3

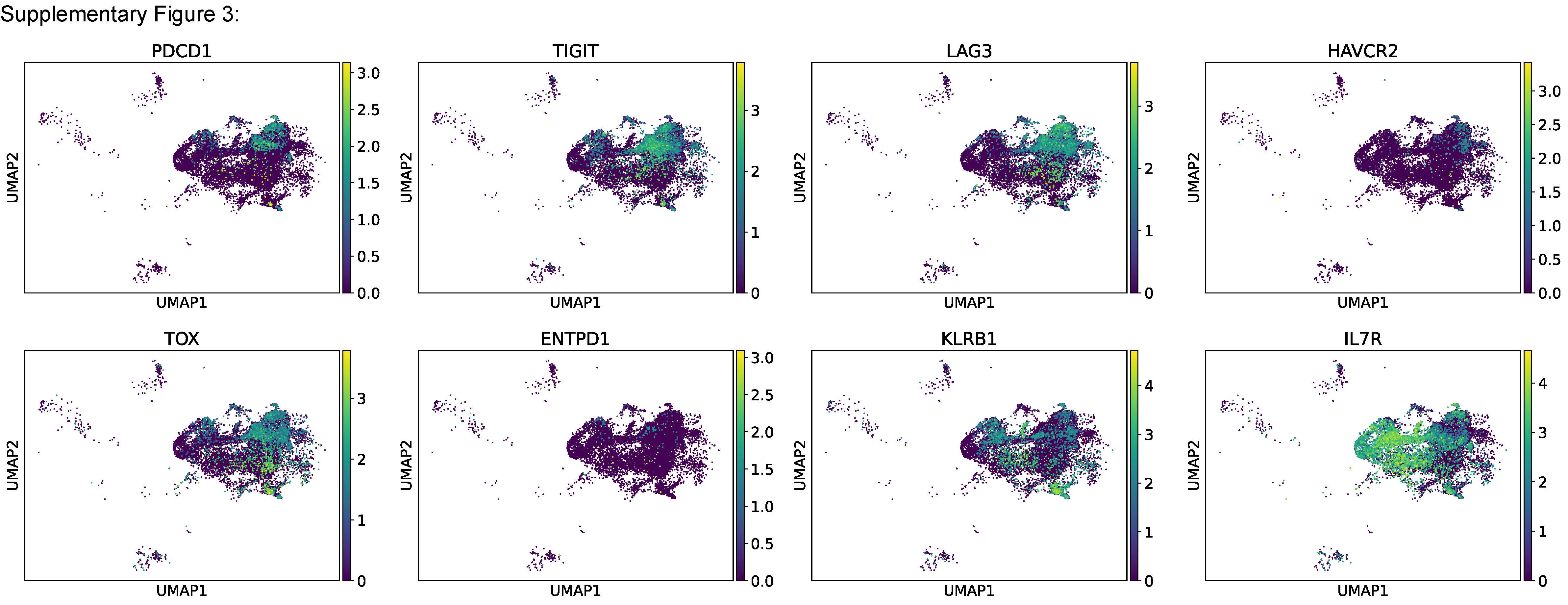
